# Voltage-sensitive dye imaging reveals inhibitory modulation of ongoing cortical activity

**DOI:** 10.1101/812008

**Authors:** Taylor H. Newton, Marwan Abdellah, Grigori Chevtchenko, Eilif B. Muller, Henry Markram

## Abstract

Voltage-sensitive dye imaging (VSDI) is a powerful technique for interrogating membrane potential dynamics in assemblies of cortical neurons, but with effective resolution limits that confound interpretation. In particular, it is unclear how VSDI signals relate to population firing rates. To address this limitation, we developed an *in silico* model of VSDI in a biologically faithful digital reconstruction of rodent neocortical microcircuitry. Using this model, we extend previous experimental observations regarding the cellular origins of VSDI, finding that the signal is driven primarily by neurons in layers 2/3 and 5. We proceed by exploring experimentally inaccessible circuit properties to show that during periods of spontaneous activity, membrane potential fluctuations are anticorrelated with population firing rates. Furthermore, we manipulate network connections to show that this effect depends on recurrent connectivity and is modulated by external input. We conclude that VSDI primarily reflects inhibitory responses to ongoing excitatory dynamics.

## Main

Electrical signaling in cortex is thought to be divided into “inputs” in the form of subthreshold synaptic potentials, and “outputs” in the form of action potentials (APs) or spikes (Grinvald and Hildesheim, 2004). Therefore, a complete understanding of cortical function requires not only a means of recording spikes in neural ensembles, but also a technique for resolving subthreshold membrane potentials (*V*_*m*_) in these populations. Voltage-sensitive dye imaging (VSDI) is a mesoscale imaging technique capable of capturing subthreshold activity across the entire rodent neocortical surface (on the order of several cm^2^) with good spatiotemporal resolution (on the order of milliseconds in time, and < 50 μm in space) (Chemla and Chavane, 2010; Ferezou et al., 2009; Grinvald and Hildesheim, 2004; Shoham et al., 1999).

In previous decades, significant progress was made in mapping the functional architecture of the brain using intrinsic optical imaging, a modality based on activity-dependent changes in the intrinsic absorptive and fluorescent properties of brain tissue. Studies combining intrinsic optical imaging with cytochrome oxidase staining revealed the interdependent organization of ocular dominance columns, cytochrome oxidase blobs (color preference), and orientation selective “pinwheel” structures in the visual system (Bartfeld and Grinvald, 1992; Blasdel, 1992a, 1992b; Frostig et al., 1990; Ts’o et al., 1990). However, intrinsic optical imaging is limited by a slow time constant (on the order of seconds (Grinvald et al., 1986, 2016)), rendering it ill-suited for capturing temporal changes in ongoing activity. To this end, VSDI has added a dynamic component to the understanding of neural assemblies. For example, VSDI-based studies have clarified the role of feedforward thalamic inputs versus intracortical recurrent activity in shaping orientation selectivity (Sharon and Grinvald, 2002), and shown how orientation-selective responses spread over the cortex as a function of stimulus shape and size (Chavane et al., 2011). VSDI has also been widely applied to the study of somatosensory computations in barrel cortex, where the somatotopic organization and spatiotemporal scale of activity is well suited to the technique. Such studies have produced important findings regarding the regulation of response dynamics by ongoing spontaneous activity (Petersen et al., 2003a), cortical state (Civillico and Contreras, 2012), behavior (Ferezou et al., 2006, 2007; Kyriakatos et al., 2017), and stimulus strength (Petersen et al., 2003b). Broadly put, VSDI has enabled the field to move beyond the static picture provided by staining and intrinsic optical imaging, adding a dynamic dimension to the understanding of mesoscale cortical organization.

However, VSDI suffers from the limitation that the superposed activity of neurites belonging to many cells is reflected in each image pixel. Uneven dye penetration, and blurring due to absorption and scattering of photons in tissue further complicate the interpretation of VSDI signals (Chemla and Chavane, 2010; Grinvald and Hildesheim, 2004; Grinvald et al., 2015). Indeed, a historical concern has been identifying which attributes of neural anatomy and physiology (e.g. layer, cell type, dendrites vs. axons, pre-vs. postsynaptic activity) are the primary drivers of VSDI measurements (Civillico and Contreras, 2006; Ferezou et al., 2006; Lippert et al., 2007; Petersen et al., 2003b). A model of VSDI that considers both the biological organization of the neuronal tissue and the physics of signal acquisition could answer these questions. However, the true power of such a model would lie not merely in inferring the relative importance of different aspects of neural activity, but also in bridging spatiotemporal scales to generate new insights into the emergent dynamics of cortical populations.

Here, we present the results of a detailed computational model of VSDI, implemented in a digital reconstruction of rodent neocortical microcircuitry (NMC), specifically, the hindlimb somatosensory cortex of a juvenile rat (Markram et al., 2015). The NMC comprises a network of ∼31,000 neurons with detailed cellular anatomy and physiology and data-driven synaptic physiology, arranged with algorithmically constrained connectivity in a 0.29 ± 0.01 mm^3^ column of tissue (Fig. 1a,b). To simulate VSDI signals, we performed simulations of the NMC to obtain *V*_*m*_ recordings of neural compartments under various experimental conditions (Fig. 1c) (see also Supplementary Fig. 2 for an evaluation of evoked VSDI in individual statistical instantiations of the NMC). Next, we corrected the compartment voltages to account for the effects of dye penetration and light transport in tissue (Fig. 1d), and collected this data into voxels (Fig. 1d,e). Using offline Monte Carlo simulations of photon-tissue interactions and a ray transfer model of microscope optics, we calculated a depth-dependent point spread function (PSF), with which we convolved horizontal slices of voxelized data (Fig. 1e,f) (see also Methods). This procedure generated a time-ordered collection of VSD images (Fig. 1g), which we repeated for various permutations of microcircuit geometry (Fig. 1h) to probe population dynamics in the NMC model.

**Fig. 1:**
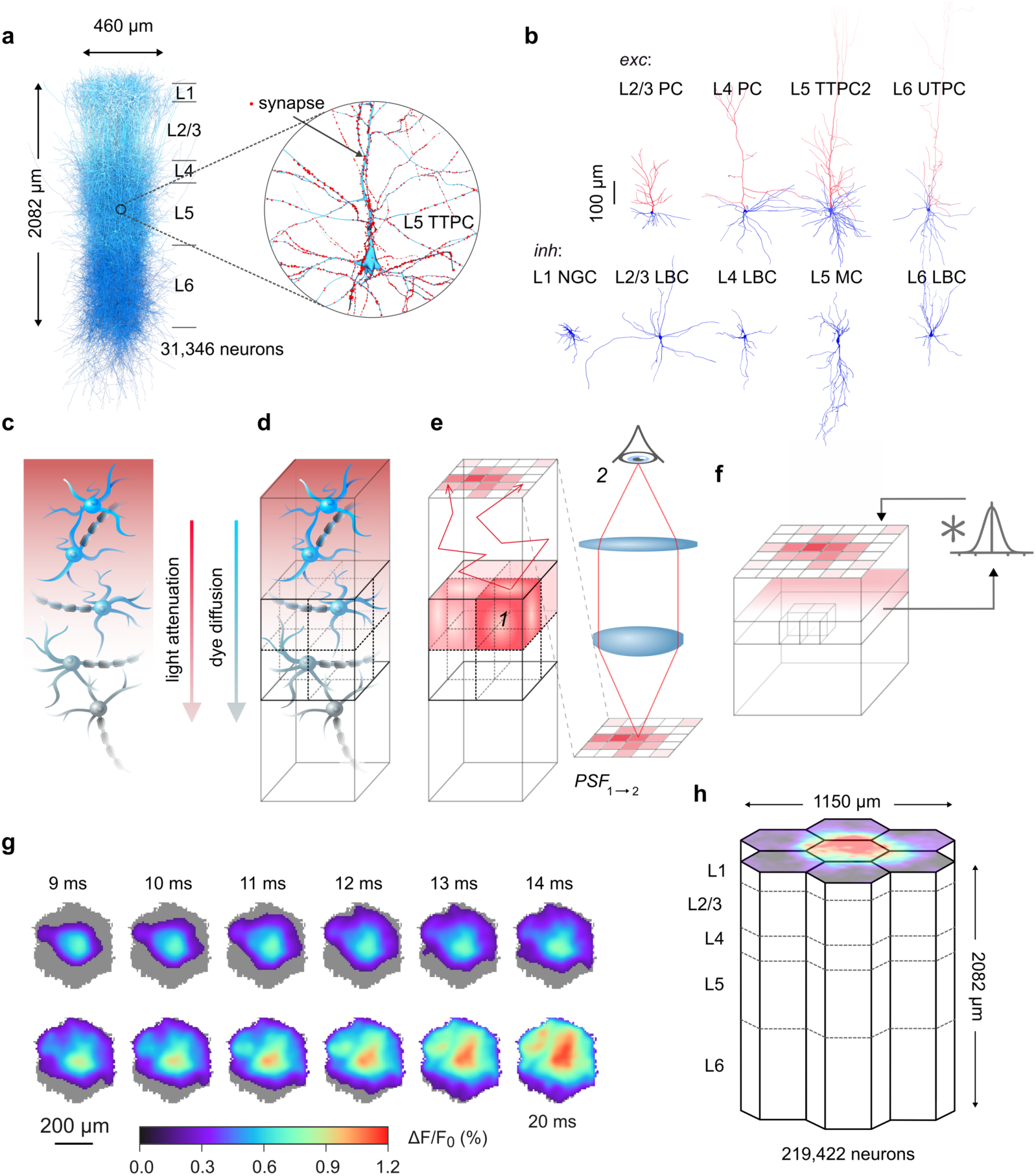
Cortical microcircuit overview and *in silico* VSDI workflow. **a**, Digital microcircuit comprising 31,346 morphologically detailed neurons connected in a columnar unit. Inset: expanded view of a L5 PC, with synapses highlighted in red. **b**, Exemplar excitatory (top) and inhibitory (bottom) cell types. Blue: axons. Red: dendrites. **c-f**, Schematic illustrating the in silico VSDI workflow. **c**, Neuron surface areas are scaled by prefactors accounting for dye diffusion and light transport in cortical tissue. **d**, Microcircuit volume is divided into voxels to facilitate calculations. **e**, Photons emitted from each voxel are scattered and absorbed throughout the tissue volume via Monte Carlo simulations. Photons reaching the cortical surface are propagated through a tandem-lens optical setup using ray transfer matrix analysis. Steps **e** and **f** are performed once for a given circuit and optical setup to determine a depth-dependent point-spread function (point-spread from 1 to 2 in panel, i.e. from voxel to camera). **f**, Raw signals at each depth are convolved with their respective point-spread function, and accumulated in a pixel array at the surface. **g**, Example VSDI image stack for 11 ms of spontaneous activity. Images were thresholded at 10% of peak response. **h**, Microcircuits were aggregated into a larger volume made of a central microcircuit column surrounding by six additional columns contacting each of the central column’s hexagonal sides (the “mosaic”). This arrangement mitigates boundary effects within the central column and facilitates the analysis of signal spread dynamics.

## Results

### Evoked VSDI response dynamics

Propagating waves of activity support the representation and integration of information in cortex (Borgdorff et al., 2007; Contreras and Llinás, 2001; Ferezou et al., 2006, 2007; Lustig et al., 2013; Petersen and Sakmann, 2001; Petersen et al., 2003b), and can be observed with VSDI. Activity spread dynamics are commonly characterized by measuring the phase velocity and aspect ratio of the propagating wavefront, the spatial extent and time-course of cortical activation, and the dependence of these quantities on stimulus strength (cf. Fehérvári et al., 2015). To quantify the similarity between the evoked response dynamics of our model and those reported in literature, we conducted a series of whisker flick-like trials and examined the spread of activity. Our stimulation protocol consisted of a single pulse of activity in 60 contiguous thalamocortical fibers emanating from a virtual ventral posteromedial nucleus (VPM) projecting to the geometric center of a concentric arrangement of 7 NMCs (the “mosaic”, Fig. 1h). We observed a radially expanding pattern of activation centered around the location of stimulus delivery, which expanded to fill the entire NMC surface over the course of several tens of milliseconds, reaching peak fluorescence at ∼57 ms poststimulus (Fig. 2a,b). The peak was immediately followed by a period of declining activity characterized by increasing hyperpolarization, which undershot baseline fluorescence, reaching a minimum at 170 ms and gradually recovering to within 10% of baseline after ∼510 ms. To quantify the temporal persistence of the signal, we calculated the half width duration (decay time to 50% of signal peak from baseline, 88 ms). The time to peak and half width duration compared favorably to the findings of Ferezou et al., 2006, who report values of ∼45 ms, and 86 ± 69 ms, respectively (Fig. 2b). We also considered the relationship between the instantaneous firing rate (3 ms bins) and the VSDI signal in a 100 ms poststimulus window (Fig. 2c,d). Our simulations indicate that peak AP firing occurred ∼7 ms *prior* to peak VSD fluorescence, contrary to the intuition that increased mean *V*_*m*_ precipitates population spiking.

**Fig. 2:**
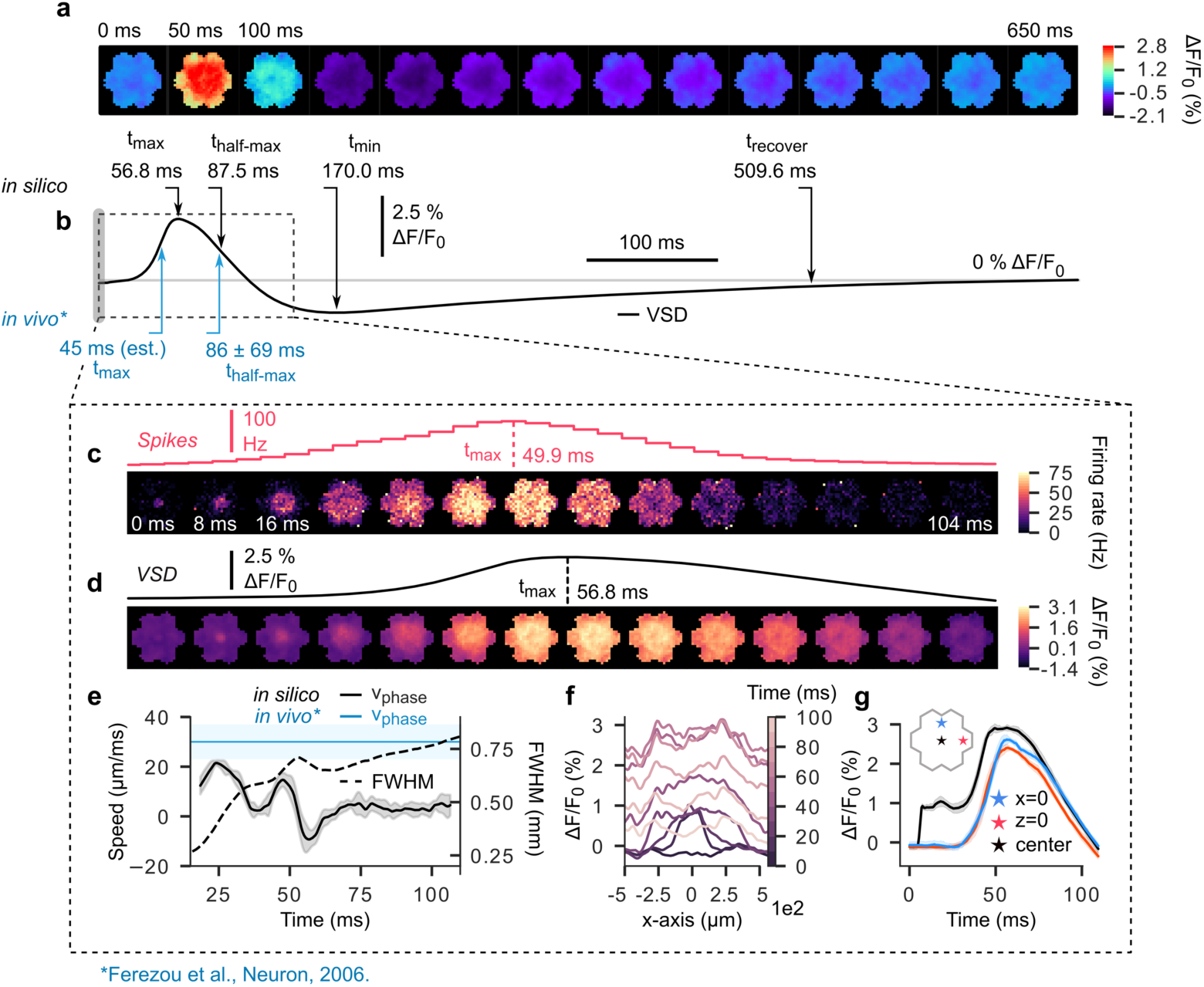
Propagation of stimulus-evoked cortical activity. **a**, 650 milliseconds of simulated stimulus-evoked VSD signals in the mosaic. The stimulus consists of a single, coincident pulse of activity in 60 contiguous thalamic projection fibers innervating the center of the interior microcircuit (delivered at t=0 ms). **b**, Time course of spatially-averaged VSD signal in **a**. Upper arrows (black) indicate points of interest along the curve. From left to right: peak latency, half-maximum duration (time to decay to 50% of signal peak), time of signal minimum, recovery time (earliest time after minimum for amplitude to stably decay to within 10% of baseline). Bottom arrows (blue) indicate in vivo values for peak latency and half-maximum duration reported in literature. **c**, Top: PSTH of spiking activity for time window (dashed box) indicated in **b** (3 ms bins). Bottom: pixel-wise PSTH (mean firing rate of all cells under each pixel) over same time window (8 ms bins). **d**,Top: expanded view of time window (dashed box) indicated in **b**, detailing ascending and descending phases of stimulus-evoked spatially-averaged VSD signal. Bottom: same as above, but for spatially extended VSD signal, where each frame was computed by averaging activity in 8 ms intervals. **e**, Activity wavefront propagation velocity (solid line, left axis), and wavefront size (full-width at half maximum, dashed line, right axis). Blue horizontal line and shaded region show in vivo measurements of stimulus-evoked wavefront propagation velocity reported in literature. Solid blue line: mean value, shaded blue region: measurement range. **f**, Spatiotemporal evolution of VSD signal along the x-axis (z=0, i.e. a horizontal line across the surface of the mosaic). Lightening hue represents passage of time. **g**, Time course of VSD activation at specific locations on the cortical surface. Blue star: circuit periphery along the y-axis. Red star: circuit periphery along the x-axis. Black star: circuit center.

Notably, a differing VSD activity time-course was observed at the stimulus delivery location relative to the NMC periphery (Fig. 2g). At peripheral points along the *x*- and *z*-axes (+540 μm and +460 μm, respectively), the rising and falling phases of the fluorescence response were almost identical to the spatial mean. In contrast, signal recorded at the stimulus location exhibited an initial transient within the first 12 ms of stimulus onset, and then gradually rose to peak fluorescence. This response pattern (initially confined, expanding thereafter) was also visible in the spatial profile of activation over time as an initially sharp peak, which gradually rose and then flattened into a plateau (Fig. 2f).

In order to characterize the propagation velocity of the evoked activity wavefront, we fit each image frame to a two-dimensional Gaussian surface and measured the change in the full width at half maximum (FWHM) over time (Fig. 2e) (see Supplementary Algorithm 1 for details). We found that activity wavefronts underwent two sequential bursts of expansion prior to peak VSD fluorescence, reaching a peak velocity of ∼20 μm/ms. Subsequently, the wavefront entered a period of contraction (−10 μm/ms) near the fluorescence peak, before gradually returning to baseline (fluctuations near zero). *In vivo* VSDI experiments have documented wavefront propagation speeds within an order of magnitude of those reported above. For example, Petersen et al., 2003a use a Gaussian fit of the cross-sectional profile of VSD images to estimate that whisker deflection-evoked waves in urethane or halothane anesthetized rodent barrel cortex propagate along barrel rows at a speed of ∼60 μm/ms, and barrel arcs at ∼33 μm/ms. See Supplementary Table 1 for a summary of cortical wavefront propagation velocities reported in literature. Also, see Supplementary Fig. 3 for an analysis of laminar VSDI activity spread in a sagittal slice (*x*-*y* plane) of the NMC.

### Excitatory neurons in layers 2/3 and 5 dominate VSDI measurements

VSDI signals are linearly proportional to the product of local *V*_*m*_ and membrane surface area (Grinvald and Hildesheim, 2004). Moreover, signals originating in neurites located in deeper layers are significantly more attenuated than those emanating from superficial layers due to uneven dye penetration and light-tissue interactions (Fig. 3b). It follows that the morphology, location, and orientation of a given cell affect the magnitude of its contribution to the optical response. To better understand these influences, we analyzed the fractional contributions of cortical layers and cell types to the overall VSDI signal. In agreement with previously reported results (Ferezou et al., 2007; Gollnick et al., 2016; Lippert et al., 2007; Petersen et al., 2003a), we found that > 90% of the raw fluorescence originated within 500 μm of the pial surface (Fig. 3b). Furthermore, we saw that neurites belonging to L2/3 and L5 neurons monopolized the “effective surface area”, which we define as the quantity that results from multiplying the original surface area of each neurite by a depth-dependent scale factor accounting for dye penetration and light transport; L2/3 and L5 contributed 44.9% and 43.7% of the total, respectively. As predicted by the distribution of effective surface area, L2/3 and L5 were the primary drivers of the VSDI signal (47.8% and 37.6%, respectively, n = 10 trials) during spontaneous activity (Fig. 3c,d). Cross-correlation revealed mutual positive correlations between the contributions of each layer and the VSDI total (Fig. 3e). However, during evoked activity, L5 contributed upwards of 67% of the signal whereas L2/3 neurites constituted 19% (Fig. 3f,g). Importantly, L5 underwent strong depolarization in the poststimulus window while L2/3 tended to hyperpolarize, indicating differential, layer-specific roles during stimulus response. Analysis of correlation supports this conclusion, showing anticorrelated activity between superficial and deep layers (Fig. 3h).

**Fig. 3:**
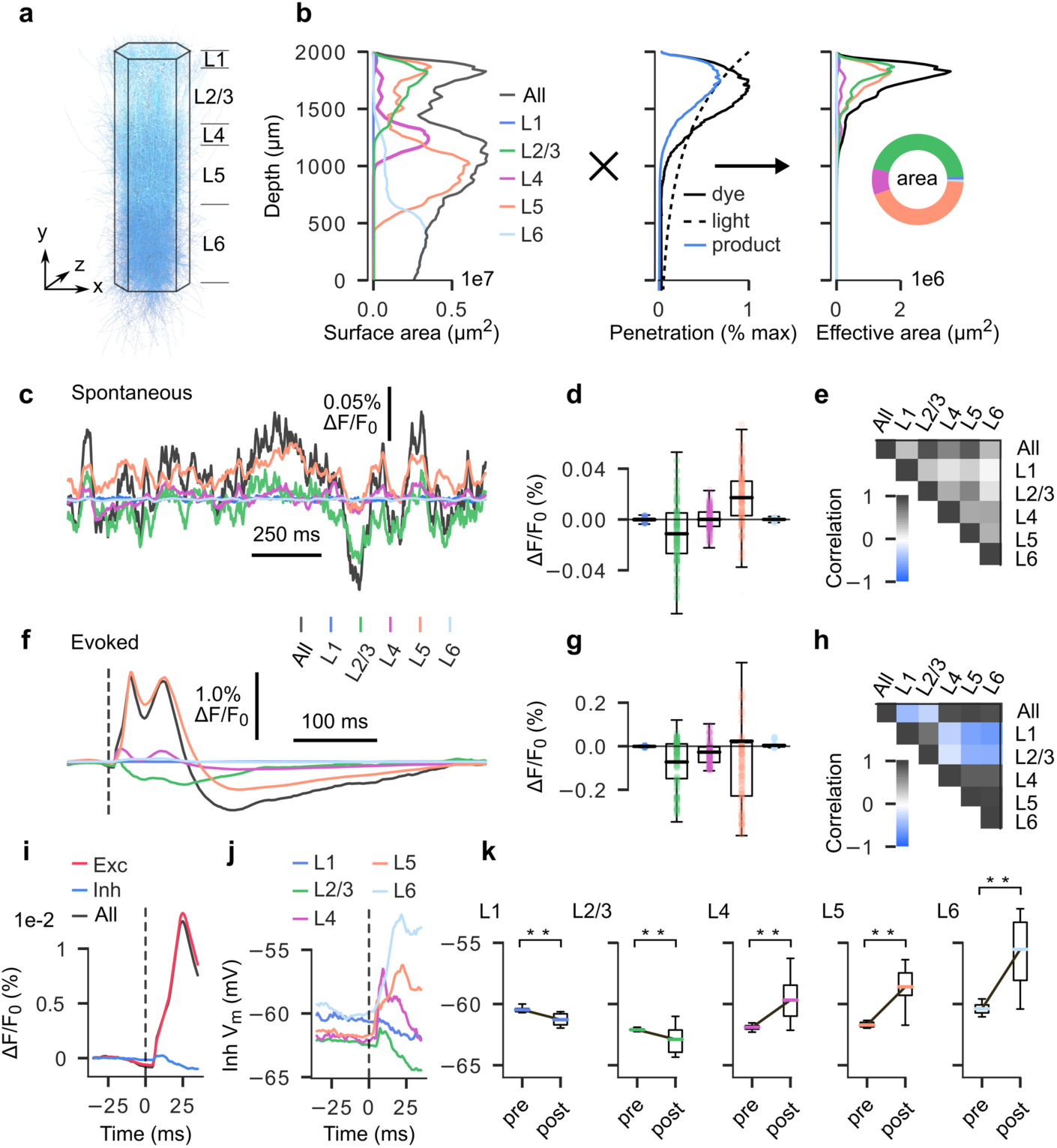
Fractional contributions to VSDI measurements by layer and cell-type. **a-b**, Surface area contributions for each layer by depth. **a**, Microcircuit, for reference respecting relative layer positions and axis orientation. **b**, (Left) Raw (unscaled) neurite surface area profiles by depth for each layer (20 µm bins). (Middle) Depth-dependent scaling prefactors accounting for dye diffusion (solid black line) and light penetration (solid blue line). Solid blue line indicates product. (Right) Effective surface area profiles by depth for each layer (raw surface area **b** scaled by product of light attenuation and dye diffusion prefactors (dashed line, middle panel), 20 µm bins). **c**, Spatially-averaged VSD signal (black) with fractional contribution of each layer (colored) for 1.5 seconds of spontaneous activity. **d**, Boxplot of fractional layer-wise contribution data in **c**, illustrating overall spread and polarity of each layer’s contributions. **e**, Correlation matrix for all traces in **d. f**, Spatially-averaged VSD signal (black) with fractional contribution of each layer (colored) for 500 ms of evoked activity. Plot begins at -50 ms, stimulus delivered at 0 ms (dashed line). **g**, Same as in **c**, but for evoked activity. **h**, Same as in **d**, but for evoked activity. **i**, Fractional contributions of excitatory and inhibitory populations to overall VSD signal, shown over a 50 ms second window spanning 25 ms pre- and 25 ms poststimulus. **j**, Mean membrane potentials computed for inhibitory cell populations in each layer, plotted over the same time window as in **i. k**, Boxplots depicting the difference between pre- and poststimulus membrane potentials for inhibitory cell populations in each layer. Boxplots in each panel were calculated using the 25 ms pre- and poststimulus periods referred to in **i** and **j**.

We also decomposed VSD fluorescence into excitatory and inhibitory components. For evoked trials, the excitatory component of the signal (> 90%) underwent large deflections in the poststimulus window, far outweighing inhibitory contributions (< 10%) (Fig. 3i). Indeed, the inhibitory fraction remained small throughout both pre- and poststimulus periods. One might expect the inhibitory VSDI fraction to increase in proportion to the excitatory fraction, as it is known that excitatory activity in healthy neocortex quickly recruits a mitigating inhibitory response, preventing runaway excitation (Fino and Yuste, 2011; Isaacson and Scanziani, 2011; Kapfer et al., 2007; Silberberg and Markram, 2007). We therefore analyzed the timecourse of mean membrane potential changes in the inhibitory populations of each layer, revealing that those in superficial layers were significantly hyperpolarized following stimulation, while those in deep layers were significantly depolarized (Fig. 3j,k). Since the dendrites of inhibitory cells tend to be spatially confined (Fig. 1b), those located in deeper layers are unlikely to contribute appreciably to the VSDI signal as their morphologies do not extend to a height reachable by dye and light. Therefore, the VSDI signal only “sees” the contributions of hyperpolarized superficial inhibitory neurons.

### Disentangling the impacts of sub- and suprathreshold neural activity on VSDI

It is thought that APs are too brief and too asynchronous to contribute substantially to the VSDI signal, despite causing large fluctuations in *V*_*m*_ (Berger et al., 2007; Civillico and Contreras, 2012; Ferezou et al., 2006; Grinvald and Hildesheim, 2004; Petersen et al., 2003a, 2003b). This conclusion is based on simultaneous VSDI and single cell patch-clamp recordings, of which spike-triggered averaging exposes the absence of individual AP waveforms from the VSDI signal (Ferezou et al., 2006). However, such experiments leave open the possibility that large *volleys* of spikes occurring within a narrow time window could still contribute to the signal. To isolate the effects of spikes on the optical response, we ran our VSDI pipeline on spike-filtered neurite compartment voltage data and compared with unfiltered data (Fig. 4a). Assuming the null hypothesis that VSDI primarily reflects subthreshold activity, we considered any difference between the raw and spike-filtered signals as “noise” due to spikes. This allowed us to calculate a signal-to-spike ratio (SSR), defined in analogy to the signal-to-noise ratio, as the squared quotient of the root mean square (RMS) amplitudes of the unfiltered signal and spiking component. That is,

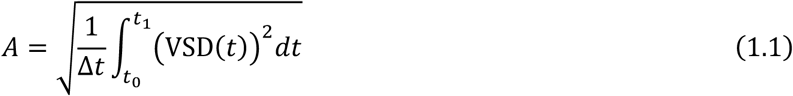

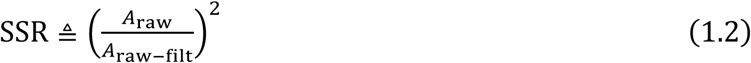

which we represented as a continuous variable by binning into 40 ms intervals with overlapping windows. Conservatively, we estimate that typical VSDI experiments have an SNR of ∼10 (Civillico and Contreras, 2005; Grinvald et al., 1999; Jin et al., 2002; Lippert et al., 2007; Tsau et al., 1996; Zhou et al., 2007). Therefore, when SSR is less than 10 (i.e., less than the empirical SNR of typical experiments), the component of the VSDI signal due to spikes is larger than contamination due to other noise sources, and in principle could be detected. Although SSR did dip slightly below our estimated detectability threshold during the poststimulus window, this is unlikely to be meaningful in most laboratory settings. However, in cases where exceptionally high SNR is achieved, information regarding the spiking component of the VSDI signal may become relevant. Therefore, we sought to understand how the frequency content of spike noise is affected by stimulation (Fig. 4b). A power spectral density analysis of spike noise immediately pre- and poststimulus showed that the frequency content differs significantly only below ∼100 Hz, with lower frequencies exhibiting greater divergence. Measurements sensitive enough to detect a spiking contribution to the VSDI signal, therefore, would only contain spike-related information below this frequency cutoff and would be dominated by low frequency components. It is important to acknowledge that (as reported previously) contributions of individual spikes are not detectable in mesoscale recordings; rather, it is the aggregate influence of population spiking that adds a small DC offset (and low frequency oscillations) to the VSDI signal, as described above. Assuming a high SNR scenario, we also undertook an analysis of the relative contributions of forward- and backward-propagating APs to the spike-related VSDI signal component (Supplementary Fig. 3). We found that nearly all of the spike-related VSDI signal is due to backward-propagating APs in dendritic arbors.

**Fig. 4:**
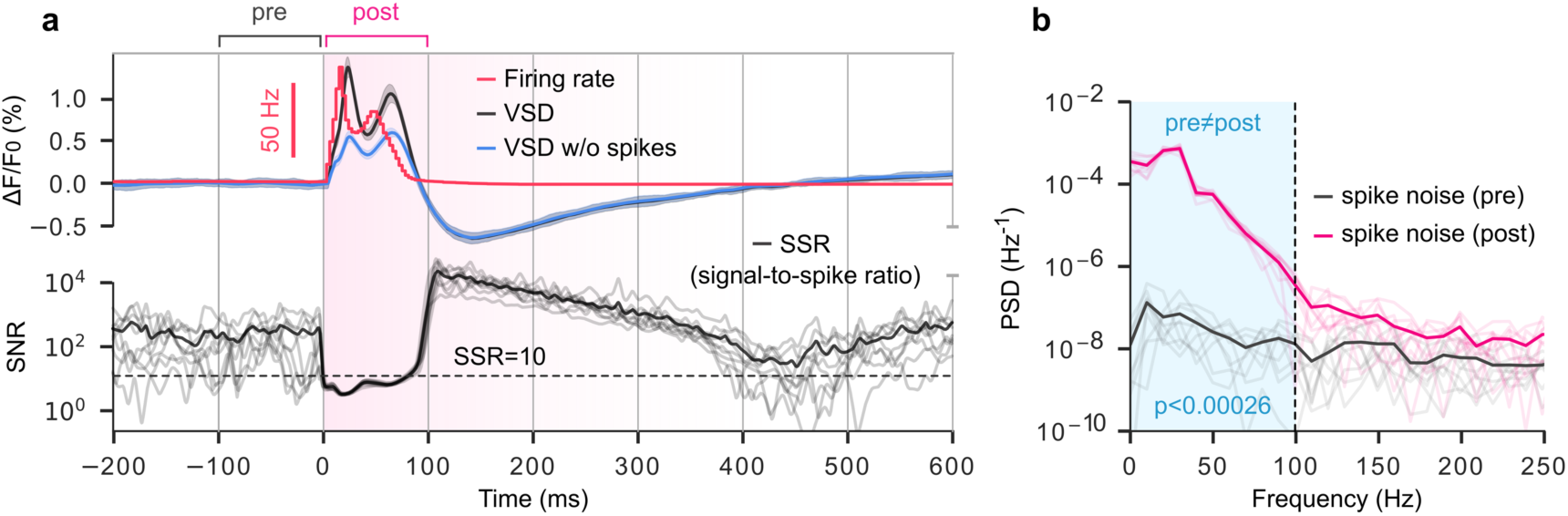
Detectability of spiking activity in the VSDI signal. **a**, Top: signal-to-noise ratio. Noise here refers to “spike noise”, or contamination of putative subthreshold measurements by APs. Dashed black line: signal-to-noise ratio = 10. Bottom: Firing rate (red), VSD (black), and VSD computed excluding spikes (membrane potentials thresholded at -55 mV, blue) for 200 ms of spontaneous activity followed by 600 ms of evoked activity. **b**, Power spectral density of spike noise computed for 100 ms prestimulus window (black), and 100 ms poststimulus window (magenta). Blue shaded box indicates frequencies for which pre- and poststimulus noise are significantly different. Dashed black line indicates frequency at which prestimulus noise and poststimulus noise are no longer meaningfully different (paired t-test, significance threshold = 0.01, adjusted to 2.6e-4 for multiple comparisons using Holm-Bonferroni correction).

### VSDI signals anticorrelate with population firing rates

In light of the unanticipated observation that spiking activity precedes VSD deflections during evoked activity (Fig. 2c,d), and the interpretation of VSDI as a measure of subthreshold *V*_*m*_, we were motivated to inspect the relationship between population firing rates and mean *V*_*m*_ in simulations of spontaneous activity. Surprisingly, this revealed a strong inverse association (Pearson correlation coefficient of - 0.86, R^2^ value of -0.83) between the two at a temporal lag of ∼23 ms, with spikes preceding *V*_*m*_ (Fig. 5a,b). The relationship between the VSDI signal and spiking exhibited a similar, albeit weaker trend, as expected in view of the correspondence between mean *V*_*m*_ and VSD fluorescence (Fig. 5c). Thus, we sought an interpretable explanation for this phenomenon in terms of population dynamics. To this end, mean-field theory was a natural choice as it provides an analytical framework for understanding the dynamics of neural populations, where each population is represented in aggregate as a single unit (Muller et al., 2007; Rudolph et al., 2004; Zerlaut et al., 2018). Since VSDI reports a summary of *V*_*m*_ across many neurons, we began by considering the relationship between mean *V*_*m*_ and conductance. Mean-field calculations for a conductance-based leaky integrate-and-fire neuron yield an “effective” membrane potential (*V*_eff_), which approximates the population mean as a function of synaptic bombardment (Rudolph et al., 2004, 2005; Zerlaut et al., 2018) (see Methods). Intuitively, *V*_eff_ may be thought of as a noisy short-term prediction of *V*_*m*_. In a hypothetical a scenario where conductances are static, *V*_eff_ is the value *V*_*m*_ converges to by exponential relaxation dynamics.

**Fig. 5:**
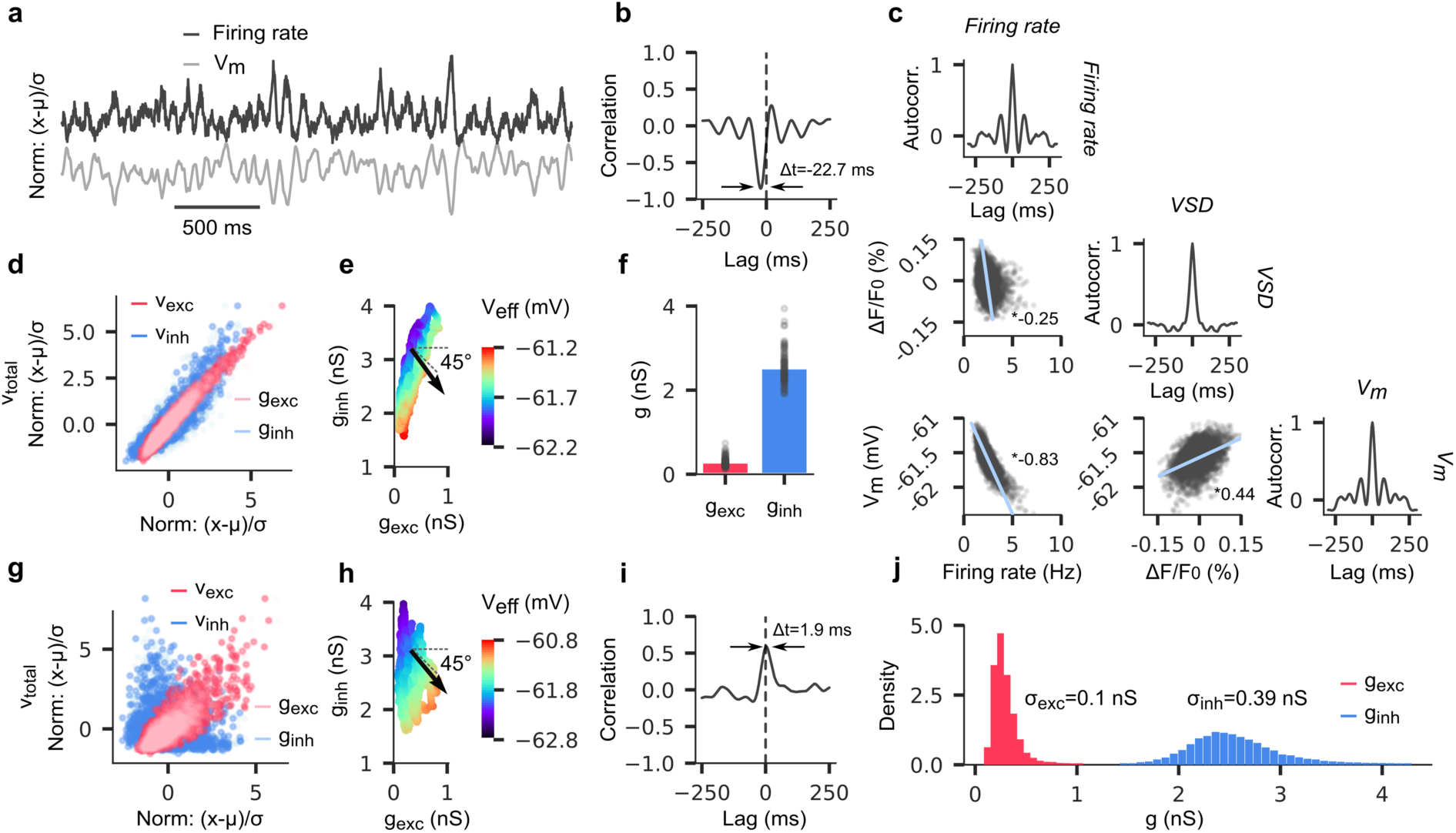
Population firing rate anticorrelates with VSDI fluctuations. **a**, Population firing rate (top, dark gray) and mean membrane potential (bottom, light gray); normalized units. Time-shifted by 22.7 ms to align signals at peak anticorrelation. **b** Cross-correlogram of population firing rate and mean membrane potential. **c**, Lower triangular scatter plot matrix depicting correlations between firing rate, VSD, and membrane potential. Starred values indicate R-squared values for linear fit. Diagonal shows autocorrelation. **d**, Scatter plot of inhibitory and excitatory conductances and firing rates (light blue, pink, blue, red, respectively) against population firing rate; normalized units. **e**, Scatter plot of excitatory vs. inhibitory conductances. Color reflects corresponding mean field-predicted membrane potential at each point in plot. Black arrow indicates directional gradient (increasing membrane potential). **f**, Bar plot of mean excitatory and inhibitory conductance values, with individual data points overlaid in black. **g**, Same as in **d**, but for decoupled network (i.e., each cell receives only pre-recorded synaptic inputs, and does not influence the network with its output). **h**, Same as in **e**, but for decoupled network. **i**, Cross-correlogram of population firing rate and mean membrane potential for decoupled network. **j**, Histogram of conductance values in **f**. Normalized so that AUC = 1 for both cell-type populations.

We proceeded in the calculation of *V*_eff_ by isolating firing rates in excitatory and inhibitory subpopulations. Experimental evidence supports the notion that excitatory and inhibitory synaptic currents are tightly balanced in healthy cortical tissue both in the resting state and during sensory processing (Denève and Machens, 2016; Sengupta et al., 2013; Zhou and Yu, 2018). Tight balance of synaptic currents implies that: 1) excitatory and inhibitory firing rates fluctuate synchronously, and 2) inhibitory conductances are greater than excitatory (Sengupta et al., 2013). Consistent with this view, we observed that excitatory and inhibitory firing rates (and thus also excitatory and inhibitory conductances) were highly correlated (Fig. 5d). Additionally, inhibitory conductances were larger than excitatory both in terms of mean and variance, with a ⟨*g*_*i*_⟩/⟨*g*_*e*_⟩ ratio of ∼9. Despite the preponderance of inhibitory conductance, we determined that *V*_eff_ was affected roughly in equal proportion per *unit* change in either conductance type (Fig. 5e). Therefore, given its much higher variance, change in inhibitory conductance was the primary driver of *V*_*m*_ fluctuations. Finally, since network balance requires high correlation between ⟨*g*_*e*_⟩ and ⟨*g*_*i*_⟩, it follows that mean *V*_*m*_, and by extension the VSDI signal, anticorrelate with total firing rate. Next, to confirm our supposition that network balance is a requirement for the aforementioned effect, we performed a series of simulations of spontaneous activity in which network connections (and thus recurrent connectivity) were disabled. Instead, each neuron was fed excitatory and inhibitory input spike trains recorded from previous simulations of the same microcircuit. Importantly, the excitatory and inhibitory inputs were derived from simulations with *different* random seeds, abolishing the coupling between excitation and inhibition (Fig. 5g), but preserving the distributions of ⟨*g*_*e*_⟩ and ⟨*g*_*i*_⟩, and therefore also their relationship to *V*_*m*_ (Fig. 5h). However, the removal of recurrent network connections inverted the correlational and causal relationship between spiking output and *V*_*m*_, as indicated by a flip in the sign of the peak correlation magnitude and its associated lag time, respectively (Fig. 5i). In the absence of network recurrence, presynaptic inputs potentiate spike firing by pushing neurons closer to AP threshold, without subsequent feedback inhibition. We conclude that VSDI primarily reflects subthreshold *V*_*m*_ changes associated with inhibitory feedback during spontaneous activity in balanced, recurrently connected cortical networks.

### Extrinsic synaptic inputs decrease the synaptic conductance ratio

In biological cortex, cellular assemblies are subject to extensive innervation by intracortical and thalamocortical projections, and by long-range projections emanating from white matter tracts (DeFelipe et al., 2002; Gil et al., 1999; Kawaguchi, 2017; Tomioka et al., 2005). Estimates of synapse counts in rat hindlimb somatosensory cortex have been reported as high as 18,000 per neuron (DeFelipe et al., 2002). Because placement of synapses in our microcircuit is constrained by the anatomical apposition of dendrites and axons (Markram et al., 2015; Reimann et al., 2015), only the formation of local synapses (∼1,145 synapses per neuron, on average) is possible, excluding significant innervation arriving from white matter. Thus, each neuron in the NMC receives a tonic injection of depolarizing current (*I*_clamp_) at its soma to compensate for missing excitatory inputs. This current is the sum of a DC component, which we compute as a percentage of rheobase (the minimal step current required to depolarize the cell to AP threshold), and a Gaussian noise component of small amplitude (Markram et al., 2015). Expressed as a percentage of rheobase, *I*_clamp_ is held constant over the entire circuit, though absolute amplitudes vary for each neuron. As a consequence, average conductance values are more than one order of magnitude smaller than those reported *in vivo* (*g*_*i*_ = 70.67 ± 45.23 nS, *g*_*e*_= 22.02 ± 37.41 nS, *g*_*i*_⁄*g*_*e*_= 14.05 ± 12.36 as measured in ketamine-xylazine anesthetized feline cortex during brainstem stimulation (Rudolph et al., 2005); cf. Fig. 5f), with concomitant reductions in variance.

We were curious to understand how the injection of current could affect the results of the previous section, and network dynamics more broadly. To this end, we attempted to replicate naturalistic conditions through the simulation of additional synaptic inputs. Assuming that most inhibitory connections occur locally (< 500 µm) (Fino and Yuste, 2011; Karnani et al., 2014; McDonald and Burkhalter, 1993), we duplicated existing excitatory synapses (5x per synapse, Fig. 6a) on a small handful of L5 PCs (n=10), and set *I*_*clamp*_ to zero. Since it is known that neurons form multisynapse connections (Deuchars et al., 1994; Frick et al., 2008; Markram et al., 1997; Silberberg and Markram, 2007; Silver et al., 2003; Wang et al., 2002), duplicated synapses for each neuron were randomly partitioned into functional groups that received identical input, representing a single connection. Group size was drawn from the distribution of synapses per connection for that cell type. Synaptic inputs consisted of spike trains generated by a homogeneous Poisson process with rate λ, which we varied incrementally until the root-mean-square error between the number of spikes in a 4 second interval in the new (added synapses) and old (somatic depolarization) simulations was minimized (Fig. 6b,c). We observed that for additional Poisson inputs at the optimal rate (0.45 Hz), mean excitatory conductance increased by a factor of ∼7, while inhibitory conductances remained largely unchanged (∼5% increase). However, conductance standard deviation changed considerably for both excitatory and inhibitory channels, increasing by a factor of 14.3 and 5.8, respectively (Fig. 6d). We also examined the relative influence of excitatory and inhibitory conductances in determining *V*_*m*_ fluctuations and found that the increased variance in *g*_*e*_ corresponded to relatively more influence than *g*_*i*_ on the trajectory of *V*_*m*_ (Fig. 6g). However, the addition of synapses did not alter the previous observation that a unit change in either *g*_*e*_ or *g*_*i*_ affected *V*_*m*_ equally. In summary, a five-fold increase in excitatory synapses with Poisson inputs in the absence of somatic depolarization tended to increase the mean of *g*_*e*_ and its influence on *V*_*m*_, and dramatically increased the variance of both *g*_*e*_ and *g*_*i*_.

**Fig. 6:**
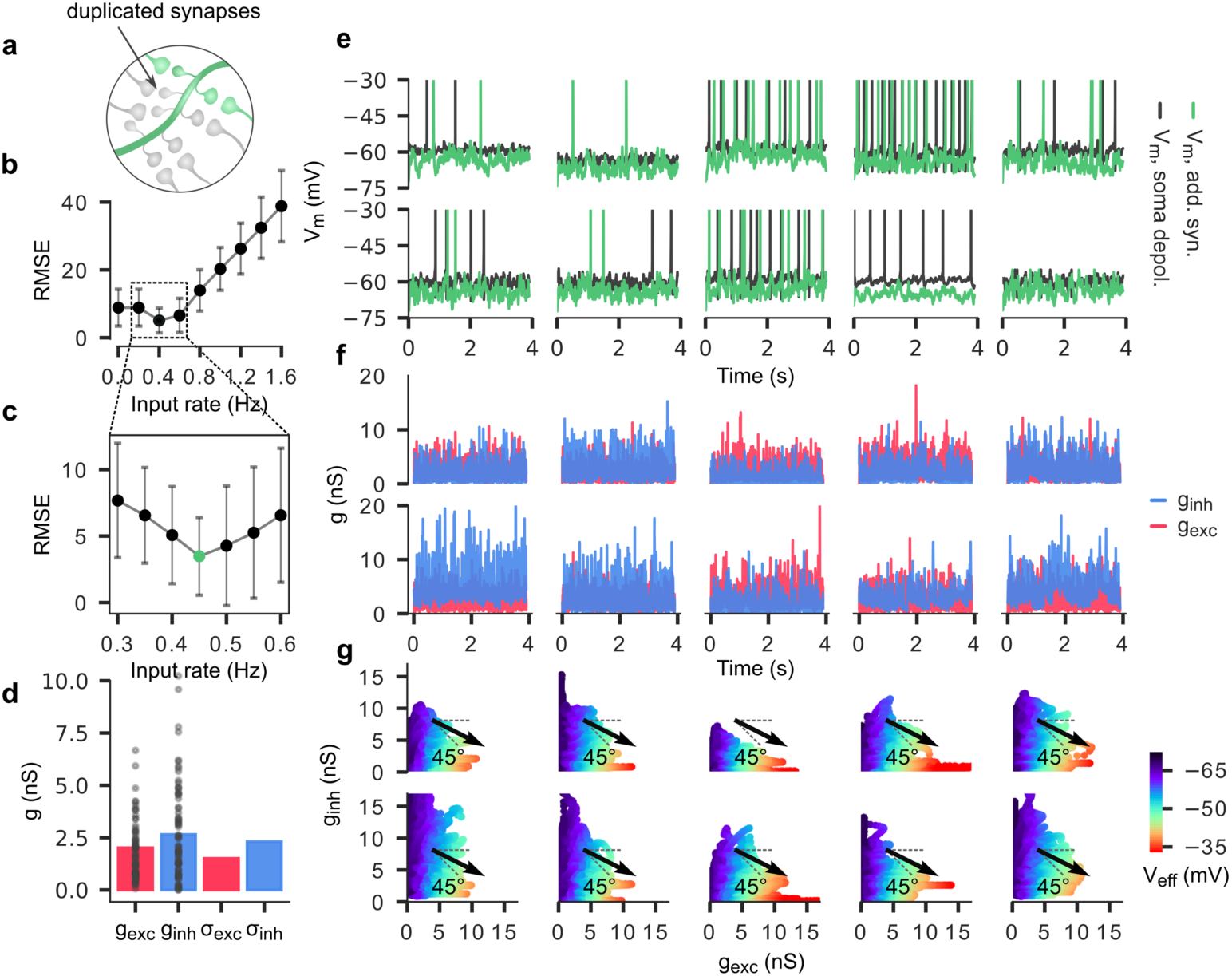
Effects of long-range excitatory synaptic inputs in the absence of somatic depolarization. **a**, Schematic showing duplication of excitatory synapses. Four identical, spatially co-located copies of each excitatory synapse were added to 10 randomly selected L5 PCs in the microcircuit. **b**, Simulated Poisson inputs were generated at different Poisson rates to evoke postsynaptic spiking. Poisson rates of the added synapses were varied to minimize the RMSE in the difference between the spiking rate of the target postsynaptic cell with and without additional synapses (but with somatic depolarization). **c**, Expanded view of **b**, for input Poisson rates between 0.3 and 0.6 Hz. Minimal RMSE indicated in green (0.45 Hz). **d**, Mean excitatory and inhibitory synaptic conductances and conductance standard deviations for all 10 neurons (pooled) for 4 seconds of spontaneous activity. **e**, Voltage trace overlays for each of the 10 neurons for 4 seconds of spontaneous activity. Green: added synapses (0.45 Hz), no somatic depolarization. Black: somatic depolarization, no additional synapses. **f**, Excitatory and inhibitory conductances (red and blue, respectively) for each of the 10 neurons for 4 seconds of spontaneous activity with added synapses at 0.45 Hz. **g**, Scatter plot of excitatory vs. inhibitory synaptic conductances. Color reflects corresponding mean field-predicted effective membrane potential at each point in plot. Black arrow indicates directional gradient (increasing membrane potential).

## Discussion

We constructed a bottom-up, biophysically detailed model of VSDI in a digital reconstruction of rodent somatosensory cortex to relate cellular anatomy and physiology to mesoscale signals, and to seek novel insights regarding cortical dynamics. As a first step, we considered VSDI measurements of evoked responses in our model and found that they were qualitatively and quantitatively similar to analogous experiments *in vivo*. Next, we used our model to deconstruct the VSDI signal into layer and cell type contributions, revealing context-dependent, strongly differentiated roles for layers 2/3 and 5. We also examined the influence of spiking activity, and found that while individual spikes are not reflected in VSDI data, large volleys of semi-synchronous spikes could affect measurements. Furthermore, our model led us to the surprising observation that the VSDI signal anticorrelates with population firing rate with a peak at a lag of ∼23 ms. Using a mean-field approach, we discovered that this is due to a predominance of inhibitory conductances, which are coupled to population spiking via recurrent connections. Finally, we considered the effects of including additional excitatory synapses to compensate for missing non-local inputs and found that a Poisson model of spiking inputs led to increased variance in values of conductance. Several details pertaining to the above results merit further discussion.

### Lateral spread dynamics of VSDI signals

As stated previously (see Results), the VSD fluorescence response to TC stimulation underwent non-uniform lateral expansion, wherein the signal quickly saturated near the location of stimulus delivery before gradually extending across the entire microcircuit (Fig. 2f,g). This observation is consistent with *in vivo* optical responses to evoked activity, in which VSDI signals saturate in a locally confined region near the stimulation site within the first 10-20 ms before expanding further (Civillico and Contreras, 2006, 2012; Fehérvári et al., 2015; Petersen, 2007; Petersen et al., 2003b). Fehérvári et al. (2015) report similar fluorescence dynamics in an *in vivo* VSDI study of mouse primary visual cortex (V1). In particular, they find that in a localized region around the site of an applied 50 μA current impulse, fluorescence rapidly increases within ∼10 ms, before saturating and then expanding laterally. They propose that the initial peak primarily reflects monosynaptic excitatory postsynaptic potentials (EPSPs), which are followed by the propagation of disynaptic activity at greater latencies. This explanation is consistent with our finding that the first response occurs locally and quickly plateaus, since it is likely due to feedforward PSPs evoked by direct TC innervation at the center of the microcircuit. Subsequent activity spread would occur only following a monosynaptic delay, as the targets of TC projections propagate the signal to their postsynaptic partners. Also, we note that in comparison to average signal transmission speeds reported in literature (Table 1), the wavefront phase velocities calculated here are relatively low. We speculate that this may be due to slicing of mid-range intracortical axons during the morphology reconstruction process. It is well known that extended axonal arbors are at risk of slicing during histological processing, and efforts were made to repair severed arbors using statistical methods (Markram et al., 2015). However, it is unlikely that such repairs would fully correct for slicing artifacts, leaving open the possibility that significant numbers of mid-range connections are missing. If true, it would tend to decrease wavefront propagation speeds, as signal transmission would be forced to proceed strictly through short-range connections.

### Lack of correlated activity between VSDI signals and layer 2/3 PCs

A point of disagreement between our results and those described in literature is the degree to which VSDI recordings are correlated with simultaneous whole-cell (WC) recordings in L2/3 (see Supplementary Fig. 1). Several *in vivo* studies have reported a high correlation between VSD fluorescence and the *V*_*m*_ of single neurons in L2/3 rodent barrel cortex (Berger et al., 2007; Ferezou et al., 2006; Petersen et al., 2003a, 2003b). However, due to the technical challenges associated with simultaneously performing VSDI and WC recordings in live animals, these studies used anesthesia (Ferezou et al., 2006; Petersen et al., 2003a, 2003b) or *in vitro* slice preparations (Berger et al., 2007) to establish the correspondence between *V*_*m*_ and VSDI traces. It has been shown that anesthetic agents increase cortical synchrony and pairwise neural correlations (Antkowiak, 2002; Greenberg et al., 2008; Kreuzer et al., 2010; Murphy et al., 2011). Of particular relevance, Greenberg et al. (2008) found that correlated AP firing in pairs and populations of L2/3 neurons in rat visual cortex increased significantly during anesthesia as compared to the awake state. Therefore, the disparity between the strength of VSDI-*V*_*m*_ correlations observed *in vivo* and those extracted from our simulations may be at least partly explained by differences in cortical state. Since VSDI signals reflect an average over *V*_*m*_ deflections in a large number of neuronal processes mostly situated in L2/3, anesthesia-induced synchrony among L2/3 neurons would tend to increase the correlation between any given L2/3 neuron and the population mean. Our model does not consider the effects of anesthesia, nor do we observe the emergence of oscillatory cortical states. Thus, both pairwise and population neural correlations remain relatively weak during spontaneous activity, resulting in a lower correspondence between VSD fluorescence and individual *V*_*m*_ measurements.

### Changes in spiking activity precede deflections in mean membrane potential

We showed that spikes precede *V*_*m*_ fluctuations during both spontaneous and evoked activity (see Fig. 2c,d; Fig. 4a; Fig. 5b), confirming several studies including one in rat barrel cortex (Petersen et al., 2003b), and two others in ferret visual cortex (Eriksson et al., 2008; Roland et al., 2006). A reasonable expectation may be that, on the contrary, increases in VSD fluorescence should precede increased spike firing, since membrane depolarization would tend to bring neurons closer to threshold making APs more likely. However, as suggested by Eriksson et al. (2008), since each cell contacts many postsynaptic partners (452 ± 272 in our microcircuit), any given AP will elicit postsynaptic potentials (PSPs) in hundreds to thousands of other cells, meaning that a mere handful of spikes can significantly impact mean *V*_*m*_ in a population. Of course, spike initiation requires membrane depolarization, but only a fraction of the population is active at once (∼26% at evoked response peak, and ∼0.4% during baseline; 2 ms bins). Therefore, *V*_*m*_ changes associated with spike firing are outweighed by downstream PSPs, with a monosynaptic delay. We found a 6.9 ms delay between peak spiking and subthreshold response to stimulation (Fig. 2c,d), and a 22.7 ms delay during spontaneous activity (Fig. 5b). Monosynaptic signal transmission reportedly requires between 6 and 14 ms in cortex (González-Burgos et al., 2000), suggesting that deflections in mean *V*_*m*_ primarily reflect monosynaptic activity in the first case (evoked), and disynaptic inhibition in the second (spontaneous). Indeed, this conclusion is supported by the reversal of sign in the correlation between *V*_*m*_ and firing rate in the putatively disynaptic, spontaneous case. Furthermore, uncoupling the network, thereby disabling recurrent disynaptic connections, abolished the temporal lag and inverse relationship between spiking and mean *V*_*m*_ (Fig. 5i), creating a situation in which increased membrane potential merely potentiates APs. Thus, VSDI may report either monosynaptic excitation or disynaptic inhibition depending on the presence or absence of external inputs. That this aspect of VSDI measurements could have been missed in previous research begs explanation. First, simultaneously performing VSDI measurements and population spike recordings is a technical challenge, and has only been attempted in a handful of studies (e.g., (Eriksson et al., 2008; Roland et al., 2006)). Moreover, the lag between VSDI signals and spike firing, in addition to any extracortical noise sources, would tend to obscure the immediate observation of a correlation. Last, VSDI signals are imperfect proxies of mean *V*_*m*_ since they are heavily biased towards contributions of neurites within L2/3 (Fig. 2b).

### Influence of cortical state on network dynamics

Regenerative activity in the microcircuit is sensitive to the concentration of extracellular calcium ([Ca^2+^]_0_) and level of tonic depolarization. As demonstrated by Markram et al. (2015), increasing tonic depolarization has the effect of pushing cells closer to AP threshold, while decreasing [Ca^2+^]_0_ (within the physiological range, 1-2 mM) tends to shift the excitatory-inhibitory balance in favor of inhibition. Varying these two parameters, they observed the emergence of four distinct regimes in the behavior of the microcircuit, characterized by the presence or absence of regenerative activity for either spontaneous or evoked conditions. Furthermore, within each regime, varying [Ca^2+^]_0_ moved the network along a spectrum between synchrony (high [Ca^2+^]_0_) and asynchrony (low [Ca^2+^]_0_). It is well established that *in vivo* concentrations of extracellular ions are maintained within relatively narrow physiological ranges by tightly regulated homeostatic pathways, and that alterations in these concentrations affect network dynamics (Barreto and Cressman, 2011; Ding et al., 2016; Gleichmann and Mattson, 2010; Henn et al., 1972; Kraio and Nicholson, 1978; Rasmussen et al., 2017). We theorize that a spectrum of network regimes similar to those observed in our network could also be present in biological cortex, and that under normal physiological conditions, the network sits at or near the transition point between regimes. This could serve to maximize sensitivity subject to the constraint of avoiding runaway excitation, thereby optimizing the potential of cortical tissue to encode sensory stimuli. Furthermore, it is known from *in vivo* recordings that cortical neurons in awake animals exhibit low input resistances, relatively depolarized membrane potentials (∼-60 mV), and significant *V*_*m*_ fluctuations (Destexhe, 2007, 2010; Destexhe et al., 2003). Collectively, these properties are referred to as the “high-conductance state”, since they are a consequence of synaptic bombardment causing mean conductances to exceed resting conductance (Destexhe, 2007). High-conductance states are thought to play an important role in determining neural response properties, with consequences for computation (Destexhe, 2007, 2010; Destexhe et al., 2003). Independent and convergent lines of evidence suggest that the critical transition point in our model (somatic depolarization at ∼100% and [Ca^2+^]_0_ = 1.25 mM) is most analogous to quiet wakefulness, with some high-conductance state properties.

Comparing the time to peak and half width duration of our evoked VSD fluorescence response to those obtained in a similar study of mouse barrel cortex (Fig. 2b) reveals a strong correspondence in the temporal profile of cortical activation for awake animals (half-width: 86 ± 69 ms), but not anesthetized (37 ± 8 ms) (Ferezou et al., 2006). Furthermore, the dominance of inhibitory conductances, relatively depolarized membrane potentials (∼-60 mV), and significant *V*_*m*_ fluctuations argue for a high-conductance-like state, which is associated with wakefulness (Destexhe, 2007, 2010; Destexhe et al., 2003). Finally, the slow-wave oscillations typically observed during sleep and anesthesia are not present in our model, though this could be a result of missing thalamocortical interactions or neuromodulation, rather than a local property of the tissue (Contreras and Steriade, 1996; Contreras et al., 1997; Mena-Segovia et al., 2008; Murphy et al., 2011; Steriade, 2000). Thus, we propose that the results of our model are best interpreted as representing an awake, *in vivo*-like state. We note at least one major caveat to the above interpretation, namely that values of synaptic conductances are significantly lower than those estimated from *in vivo* recordings in the awake state. Since magnitude of conductance in dendritic arbors is known to affect the integrative properties of neurons (Destexhe, 2007; Destexhe et al., 2003), this discrepancy is likely to affect network dynamics. However, we leave a detailed analysis of the consequences for future research.

### VSDI-firing rate anticorrelation lag time: an index of locality?

We have shown that during spontaneous activity, population firing rate anticorrelates with mean *V*_*m*_ and spatially averaged VSDI data for recurrently connected balanced networks. However, this observation was made in the context of an isolated local network, suggesting that a relaxation of these conditions could moderate the effect. To this end, we attempted to replicate the effects of extrinsic (non-local) synaptic inputs and assessed changes in the mean, standard deviation, and ratio of *g*_*e*_ and *g*_*i*_. Due to resource constraints, we were restricted to NMC simulations with additional extrinsic inputs at a mere handful of synapses in the network, and were thus unable to evaluate alterations to the relationship between mean *V*_*m*_ and mean firing rate as a function of long-range input statistics. However, the preliminary results indicate a shift in the mean conductance ratio in individual neurons, suggesting that extrinsic inputs can influence local network properties. Indeed, Roland (2017) suggests that network balance is a local property that is tightly maintained during spontaneous activity, but which may be temporarily disrupted by a quick succession of excitatory APs from non-local cortical regions. For sufficiently powerful inputs, local inhibitory neurons cannot fully compensate for increased excitatory activity, and additional inhibitory cells must be recruited to prevent runaway excitation (Dehghani et al., 2016; Huys et al., 2016; Roland, 2017). The process of recruiting additional inhibition may introduce a delay between peaks in inhibitory and excitatory firing, relaxing the tight balance of activity in the network. Thus, we propose that the inverse relationship between VSDI signals and mean firing rate revealed by our model could provide an “index of locality”, i.e. an increased delay in the peak anticorrelation between VSDI and population spiking represents a stronger perturbing extrinsic influence. Mechanistically, the delay between changes in firing rate and VSDI fluctuations reflects the time required after a spike is fired for recurrently connected inhibitory neurons to respond by hyperpolarizing the *V*_*m*_ of the spiking cells. We predict that stronger peripheral inputs or corticocortical interactions will result in greater lag times for peak anticorrelation between VSD fluorescence and population firing rates. Indeed, a preliminary analysis of the evoked response to TC stimulation (see Supplementary Fig. 5) revealed a significant shift in the anticorrelation lag time as compared with spontaneous activity (23 ms vs. 130 ms), consistent with this prediction. Currently, the bulk of evidence for network balance has been drawn from correlations in the membrane potentials of nearby cells with similar orientation tuning, and measurements of conductance ratios over time in individual cells (Denève and Machens, 2016). Our proposal for the novel use of VSDI data to determine a locality index would add confirmatory evidence for the balance of inhibition and excitation at the *network* level, while simultaneously providing a metric for evaluating the degree of influence of non-local activity on local microcircuitry. We leave a thorough exploration of this hypothesis and its implications for future studies.

### Limitations and outlook

As regards the validity of the *in silico* model of VSDI presented here, we propose the consideration of three conceptually distinct layers: 1) whether the biophysical model of VSDI, i.e. the calculations linking cellular activity to measured fluorescence, reasonably approximate reality, 2) whether the composition and architecture of the tissue itself is biologically plausible, 3) whether the simulations are *functionally* representative of biological neocortex.

On the first account, we assert that the excellent linearity and fast kinetics of VSDs (Lippert et al., 2007) greatly simplify their analytical relationship to *V*_*m*_. Regarding the second concern, a caveat is the absence of several important structural details from the version of the NMC used here, including glial cells, vasculature, and long-range intracortical axons (Markram et al., 2015). However, in the case of glia, the slow timescale of response (3-4 ms (Schummers et al., 2008)) and small amplitude of *V*_*m*_ deflections (1-7 mV (Kelly and Essen, 1974)) make it unlikely that they contribute meaningfully to VSDI signals. As regards vasculature, since our *in silico* VSDI pipeline already accounts for the bulk optical properties of cortical tissue (see Methods), their effects have, in principle, been accounted for. With respect to the third concern above, several details that are likely to influence network dynamics, including gap junctions, multivesicular release, neuromodulation, and synaptic plasticity, are not present in this version of the NMC.

In addition, we note several caveats concerning the comparison of our *in silico* stimulation protocol to whisker deflection experiments in rodents. First, as a model of hindlimb somatosensory cortex, the NMC lacks the unique cytoarchitecture and anatomical organization that characterize barrel cortex (Schlaggar and O’Leary, 1994; Simons et al., 1984). Furthermore, the NMC excludes the trigeminal and thalamic nuclei, and therefore does not exhibit sensory processing delays (and VSDI response latencies) or the modulation of cortical dynamics by TC feedback. Previous experiments have shown that cortical activation patterns depend on stimulus strength, with a tendency for excitation evoked by weak stimuli to remain confined to a single barrel (Berger et al., 2007; Fehérvári et al., 2015; Gollnick et al., 2016; Petersen et al., 2003b). In our model, TC projection fibers innervating the geometrical center of the microcircuit fire a simultaneous AP, a construction that does not capture the full complexity of afferent TC signaling nor permit modification of the stimulus strength in a biologically plausible way. Finally, experiments have implicated reciprocal TC pathways (Bazhenov et al., 2002; Hughes et al., 2002; Steriade et al., 1993) and intracortical interactions (Timofeev et al., 2000) in the emergence of slow wave activity. It is known that cortical oscillations interact with sensory responses to produce differentiated VSDI signals (Petersen et al., 2003a). Thus, an *in silico* account of the effects of brain state on VSDI measurements awaits future iterations of the NMC that include TC feedback and corticocortical interactions.

In view of these concerns, future versions of BBP brain models that extend well beyond the microcircuit scale are forthcoming. An anatomically complete reconstruction of the somatosensory cortex (and ultimately the entire neocortex) featuring biologically appropriate macro- and micro-connectivity (Reimann et al., 2019) will help to resolve questions concerning the effects of missing long-range inputs. Additionally, a model of neuromodulatory dynamics, the effects of which are known to be implicated in the transitions between, and maintenance of, cortical states (Colangelo et al., 2019), will support the investigation of state-dependent VSDI responses. Finally, a detailed model of the neuro-glia-vasculature ensemble will pave the way for future simulation-based studies of imaging techniques such as fMRI that rely on blood-oxygen-level-dependent (BOLD) signals.

### Concluding remarks

This study demonstrates the utility of bottom-up biophysical modeling as a complement to experimental approaches for understanding the relationships between spatial and temporal scales of cortical signaling. Using our model, we clarified which aspects of neural anatomy and physiology shape VSDI signals. Additionally, we discovered that during ongoing spontaneous activity, VSDI primarily reflects subthreshold activity associated with recurrent inhibitory connections. These insights were gleaned from *in silico* data beyond the reach of current experimental techniques.

## Methods

### Microcircuit

Our *in silico* VSDI model was implemented in a digital microcircuit consisting of a connected network of 31,346 neurons, ∼8 million connections, and ∼37 million synapses. The network was arranged in a columnar volume 462 × 400 µm wide, and 2082 µm deep. A spatially extended version was constructed by interconnecting 7 such units in a hexagonal tiling (the “mosaic”). In this configuration, depth axes were mutually parallel, and columnar surfaces were coplanar (Fig. 1h). The cell morphologies populating the circuit were obtained from 3D reconstructions of biocytin-stained neurons from juvenile rat hindlimb somatosensory cortex, while the placement, connectivity, and electrophysiological properties of each cell was determined algorithmically and constrained by sparse data derived from experiments and literature (Markram et al., 2015). TC innervation was reconstructed using VPM axon bouton density profiles in rat barrel cortex, and synapses were probabilistically assigned to incoming fibers using a Gaussian distribution centered around each fiber (Markram et al., 2015). See Markram et al. (2015) for additional details concerning microcircuit construction and composition.

### Supercomputing

A 2-rack Intel supercomputer using dual socket, 2.3GHz, 18 core Xeon SkyLake 6140 CPUs, with a total of 120 nodes, 348 GB of memory, and 46 TB of DRAM was used to run the simulations and carry out analysis.

### Simulation

Simulations were conducted using proprietary software based on the NEURON simulation environment (Hines and Carnevale, 1997). Data were output in the form of binary files containing spike times and compartment *V*_*m*_ sampled every 0.1 ms for each neuron in the network. Extracellular calcium and potassium concentrations, as well as somatic depolarization are free parameters in the model, and were set to 1.25 mM, 5.0 mM, and ∼100% threshold, respectively, to most closely mimic an in vivo-like network state (Markram et al., 2015).

Trials simulating evoked responses modeled TC stimulation with of a single pulse of activity in 60 contiguous thalamic fibers projecting to the geometric center of the microcircuit. For experiments requiring a larger cortical surface area, the same approach was applied to the spatially extended hexagonal microcircuit tiling (see Microcircuit above). Activity was simulated over 10 trials (i.e. random seeds) for a duration of 5 seconds each, with the first 2 seconds of data in each trial discarded to avoid any boundary condition-dependent artefacts. The stimulus was delivered at 2500 ms, meaning that each trial consisted of an initial period of 500 ms of spontaneous activity, followed by 2500 ms of poststimulus activity.

### Signal calculation

We assumed that the VSDI signal emanating from a small patch of cellular membrane was linearly related to the product of the membrane surface area and *V*_*m*_ (Berger et al., 2007; Ferezou et al., 2006, 2009; Gollnick et al., 2016; Grinvald and Hildesheim, 2004; Lippert et al., 2007; Petersen et al., 2003a, 2003b). Our neuronal morphologies are composed of small segments (“compartments”) of equipotential cable whose surface area and transmembrane voltage were multiplied to obtain the raw VSDI signal. This raw signal was scaled for each compartment as a function of depth to account for the physics of dye diffusion and the scattering and absorption of illumination light (Fig. 1d). The degree of signal attenuation due to uneven staining through the cortical depth was interpolated from data measured in four mouse brains treated with RH1691 voltage-sensitive dye, flash-frozen and sliced into 20 μm thick cryosections (Ferezou et al., 2006). To reduce data storage requirements, we divided the microcircuit into voxels, within which an aggregate signal was computed by summing the contributions of all compartments in that voxel:

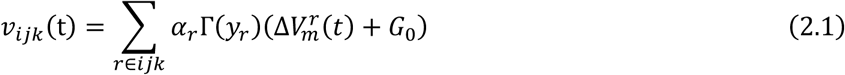

where *ν*_*ijk*_ denotes the value in the *ijk*^th^ voxel, *α*_*r*_ is the surface area of the *r*^th^ compartment, Γ(*y*_*r*_) is an attenuation prefactor accounting for dye penetration and scattering/absorption of illumination light at depth *y* for compartment *r*, 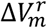 is the change in membrane potential for the *r*^th^ compartment, and *G*_0_ is a constant reflecting the combined reflecting the combined contributions of background noise and autofluorescence (assumed isotropic). The value of *G*_0_ was fixed by requiring that a 10 mV change in *V*_*m*_ correspond to a ∼0.5% change in fluorescence over baseline (ΔF/F_0_), as reported by Ferezou et al. (2006), assuming an average resting potential of -65 mV.

To model the effects of scattering and absorption in the tissue, we used a Monte Carlo simulation-based approach (see Point Spread Function) to compute an effective point spread function (PSF) for increasing depths along an axis perpendicular to the cortical surface. We used the PSF at each depth to determine the standard deviation of a Gaussian kernel, which we convolved with the horizontal data slice at that depth:

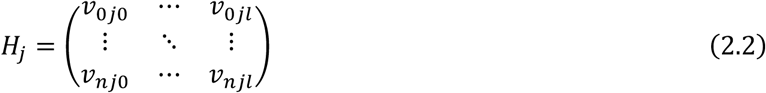

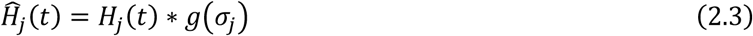

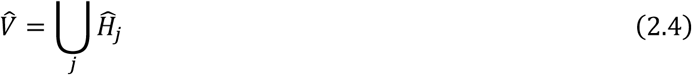

*H*_*j*_ (Equation. (2.2)) is a horizontal data slice at depth *j*, where *i* ∈ {0, …, n} and *k* ∈ {0, …, *l*}. In Equation (2.3), 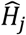 is the filtered data slice at depth *j, H*_*j*_ is the original data slice, and *g* is a Gaussian kernel, with depth-dependent standard deviation *σ*_*j*_. The union of all filtered slices yields the filtered data volume 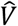 (Equation (2.4)). Post-convolution, each vertical (*j*-axis) column of voxels was accumulated into a single value, resulting in a two-dimensional matrix of pixels, which we stored as an image (Equation (2.5)). VSDI signals were computed as a fractional change in fluorescence over resting intensity (Ferezou et al., 2007, 2009; Kleinfeld and Delaney, 1996; Orbach et al., 1985; Shoham et al., 1999). This gives raw and normalized signal intensities for each pixel in an image matrix:

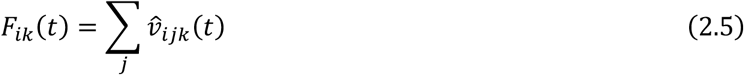

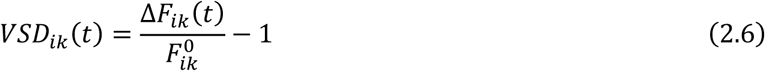

where 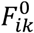 is a baseline fluorescence image obtained by averaging the first 100 frames (50 ms of data sampled at 2000 Hz). We used Equation (2.6) to calculate voltage-sensitive dye signals in this work.

### Point spread function

We calculated an empirical, depth-dependent point spread function (PSF) to account for blurring in the final image due to both scattering of emitted fluorescence photons in cortical gray matter, and also optical distortions caused by out-of-plane signal. Our method for calculating the PSF consisted of two steps: first, we used a Monte Carlo (MC) simulation-based approach to model the scattering and absorption of photons emitted from a point source within the tissue volume, varying the depth of the source in 50 micron increments; second, we used ray transfer matrix analysis to trace the trajectories of these photons through a tandem-lens optical system onto a sensor at the image plane.

MC simulations were carried out using a proprietary library built on an open-source framework for physical rendering using backward MC ray tracing, the Physically-based Rendering Toolkit (PBRT) (Pharr et al., 2016). We extended the PBRT framework to simulate photon interactions with highly turbid media using forward MC simulations based on an algorithm proposed by Abdellah et al. (2017). To determine the PSF, we moved an isotropically radiating point source of 10^8^ photons throughout a semi-infinite (lateral extent) volume of tissue, beginning at the bottom of the microcircuit in increasing increments of 50 μm, allowing each photon to scatter until it was either absorbed, or exited the cortical surface. Coefficients of reduced scattering and absorption at ∼665 nm were taken to be 4 mm^-1^ and 0.4 mm^-1^, respectively, interpolated from optical measurements made in rat gray matter for wavelengths of light spanning 450 to 700 nm (Mesradi et al., 2013). We chose the wavelength to correspond to peak emissions in the RH-1691/1692 family of blue voltage-sensitive dyes (Berger et al., 2007; Ferezou et al., 2006; Petersen et al., 2003a; Shoham et al., 1999). Refraction at the tissue-air interface was calculated using the vector formulation of Snell’s law. Using ray transfer matrix analysis, photons emanating from the tissue surface were propagated through an optical system modeled after a tandem-lens epifluorescence macroscope setup first proposed by Ratzlaff and Grinvald (1991), and subsequently used in several VSDI studies (Ferezou et al., 2006; Petersen et al., 2003a, 2003b). The system consists of two compound lenses (modeled using the thin lens approximation) set to infinite focus and placed face-to-face (Ratzlaff and Grinvald, 1991). Optical parameters (focal length, f-number and working distance) were taken from Petersen et al. (2003b), resulting in a focal plane ∼300 μm below the pia. The point source produced a sunburst image pattern on the detector array for each depth, to which a two-dimensional Gaussian surface was fit using a non-linear optimizer (Python). From these surfaces, we extracted the average spatial standard deviation, and fit the resulting array of values to a decaying exponential function to determine a depth-dependent PSF for the entire tissue-lens system. The standard deviations extracted from our PSF were used to calculate spatial kernel widths for convolution of the data with a Gaussian filter (see Equation (2.3)).

### Mean-field equations

Following the approach of Dorn and Ringach (2003), Kuhn et al. (2004), Muller et al. (2007), and Zerlaut et al. (2018), the membrane potential of a LIF neuron evolves according to:

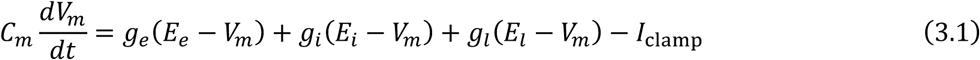

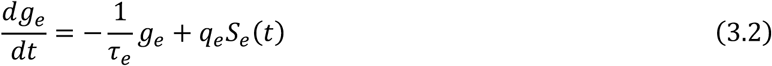

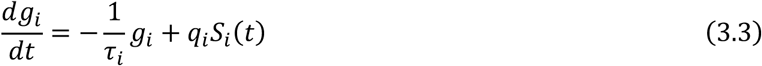

where *C*_*m*_ is membrane capacitance, *g*_*e*_, *g*_*i*_, and *g*_*l*_ are excitatory (AMPA and NMDA), inhibitory (GABA) synaptic conductance, and leak conductance, respectively, *E*_*e*_, *E*_*i*_, and *E*_*l*_ are the respective reversal potentials, *τ*_*e*_ and *τ*_*i*_ are the excitatory and inhibitory time constants, *q*_*e*_ and *q*_*i*_ are the quantal synaptic conductance increases, and *S*_*e*_ and *S*_*i*_ are presynaptic spike trains (Dorn and Ringach, 2003; Kuhn et al., 2004; Muller et al., 2007; Zerlaut et al., 2018). The term *I*_*clamp*_ represents a depolarizing current injected at the soma to compensate for missing external inputs to our microcircuit (see Results). Defining a new quantity, the “effective potential”, as:

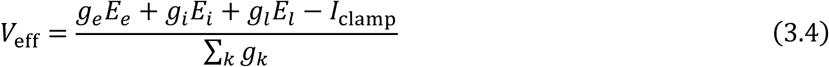

allows us to rewrite Equation (3.1) in terms of the true and effective potentials, and rearrange to obtain:

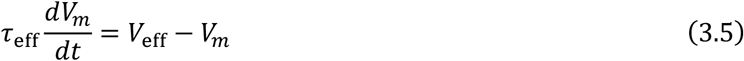

where

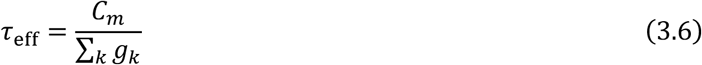

We note that Equation (3.5) has the form of an RC low-pass filter, with a cutoff frequency given by 1*/τ*_eff_; therefore we expect *V*_eff_ to be a reasonable approximation of *V*_*m*_, provided the frequency of characteristic fluctuations in *V*_eff_ don’t exceed 1*/τ*_eff_. The above equations hold true both for a single neuron, and in expectation across a population of neurons. Furthermore, if *S*_*e*_ and *S*_*i*_ are inhomogeneous Poisson processes, then in expectation, the terms *S*_*e*_ and *S*_*i*_ in Equations (3.2) and (3.3) are replaced with time-varying mean firing rates (Muller et al., 2007). Thus, in expectation,

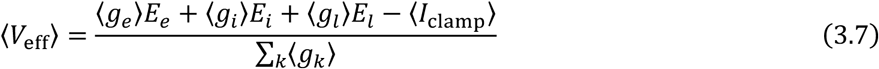

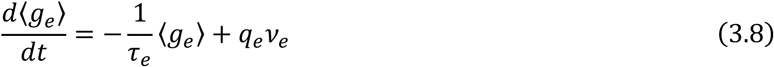

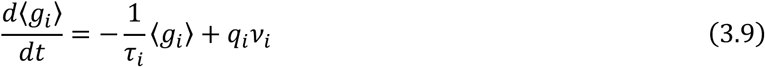

Numerical integration of Equations (3.8) and (3.9) allowed us to analytically relate mean firing rates to *V*_*m*_.

### Post-processing and analysis

All code for analysis was written in the Python programming language.

## Acknowledgements

We thank M. Nolte, M. Reimann, G. Chindemi, S. Ramaswamy, C. Colangelo, and other members of the Blue Brain Project for useful feedback and discussions, and C. Favreau for visualization support. The authors are also grateful to J. King, M. Gevaert, and W. Van Geit for technical assistance. This study was supported by funding to the Blue Brain Project, a research center of the École polytechnique fédérale de Lausanne, from the Swiss government’s ETH Board of the Swiss Federal Institutes of Technology.

## Author information

### Contributions

T.H.N., E.B.M., and H.M. conceptualized the study. T.H.N. designed the experiments, analyzed the data, prepared the figures, and wrote the manuscript. M.A. performed the Monte Carlo simulations of photons in brain tissue. G.C. developed the software for voxel-based calculation of the VSDI signal from membrane voltage data in consultation with T.H.N.

