## Supplementary information for "Voltage-sensitive dye imaging reveals inhibitory modulation of ongoing cortical activity"

### Fig. 1: Pairwise $V_m$ correlations

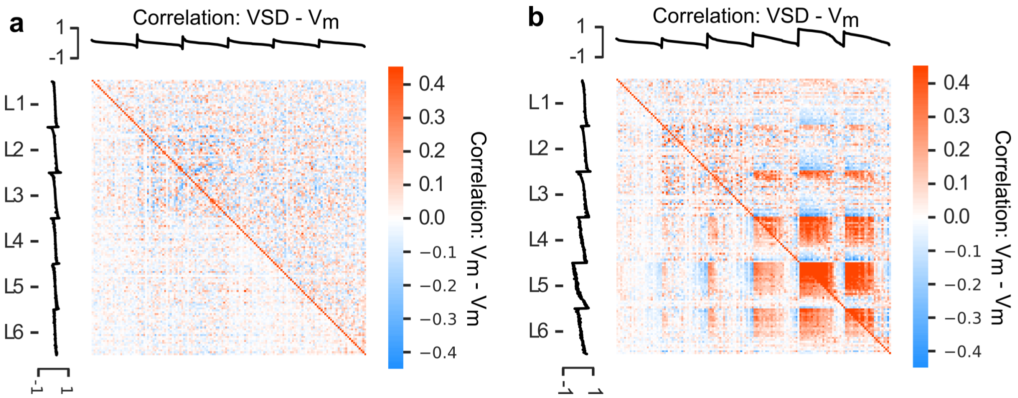


**a**, Pairwise membrane potential correlations between neurons (300 per layer) for spontaneous network activity. Upper triangle: correlations computed using thresholded traces (-55 mV). Lower triangle: correlations computed on raw traces including spikes. Top and left margins: $V_m$-VSD correlations for each cell, sorted by strength within each layer (filtered and unfiltered, respectively). **b**, Same as in **a**, but for evoked activity (single stimulus, 60 contiguous TC fibers at NMC center).

### Fig. 2: Individual NMC responses

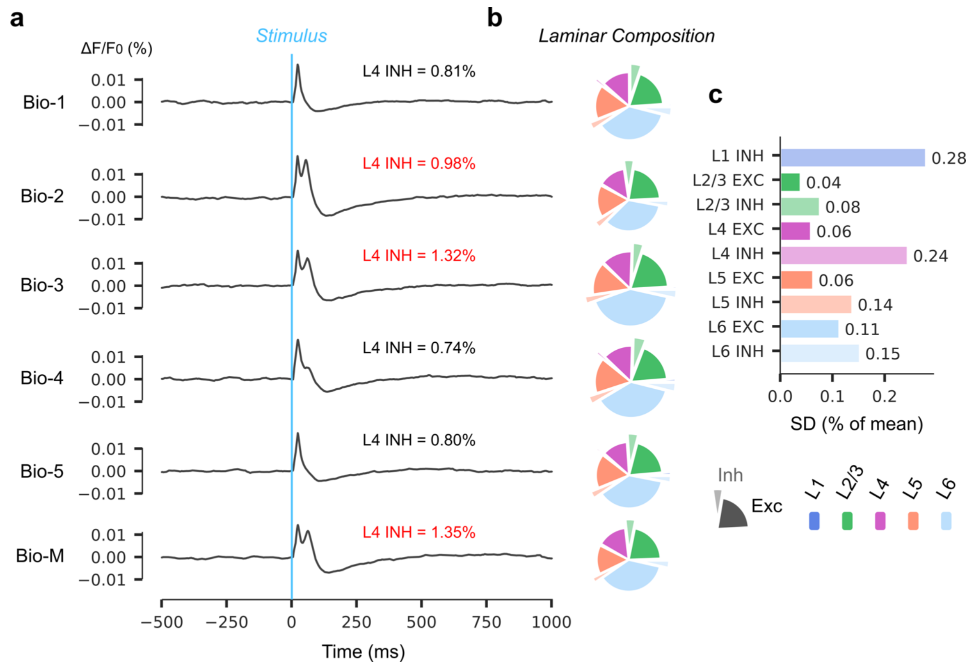


**a**, Spatially averaged VSD data for Bio-1-5 and Bio-M, $[\mathrm{Ca}^{2+}]_o = 1.25\ \mathrm{mM}$. Vertical blue line indicates stimulus. Text indicates percent deviation from mean in L4 inhibitory populations (black: subcritical response; red: supercritical response). **b**, Number of cells by layer (color) and cell type (standard or exploded pie slices) for each individual microcircuit. Radius of each pie plot is proportional to the total number of neurons. **c**, Standard deviations (percent deviation from mean) for numbers of neurons by layer and cell type.

### Fig. 3: Sagittal view VSDI dynamics

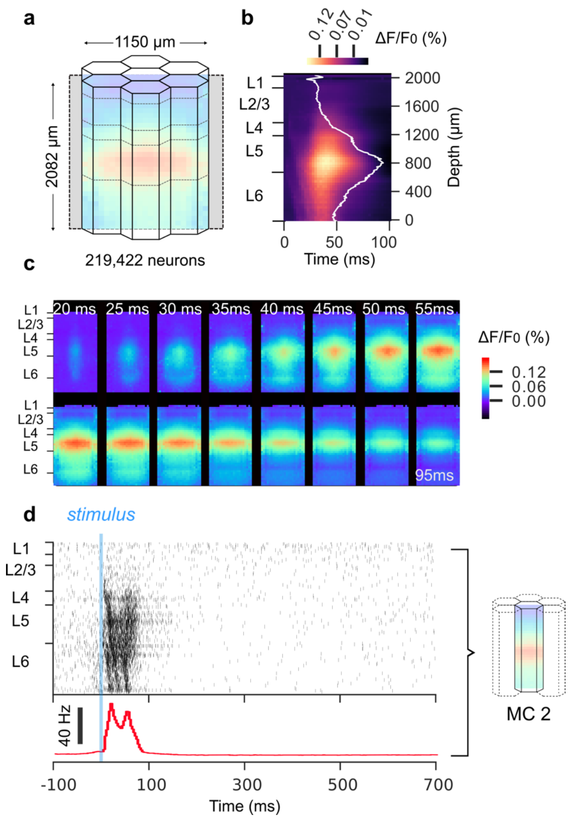


**a**, Mosaic configuration of NMC model (7 concentric columns), with sagittal imaging plane (indicated in gray) bisecting the volume along the y-axis (depth). **b**, Linescan of cross-sectional VSD activity: matrix of time series data for a vertical line through the center of the imaging plane. White line overlay is sum of each matrix row (i.e. the integral over time for each depth). **c**, VSDI data for a sagittal slice through the mosaic in 5 ms intervals (20-95 ms post-stimulus). **d**, Top: raster plot of 2000 randomly sampled cells in the central column (MC2). Bottom: same data as above, but in time histogram format.

### Fig. 4: Forward- and backward-propagating action potentials

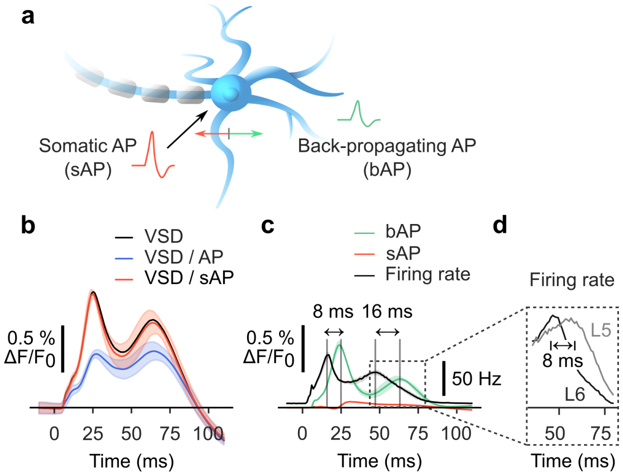


**a**, Schematic illustrating forward- and backward-propagating APs. Red: APs occurring at the soma/axon initial segment (sAP). Green: back-propagating APs occurring in dendritic arbors (bAP). **b**, Time domain comparison of filtered VSD signals. Black: full signal, no filtration. Blue: VSD signal computed with thresholded $V_m$ (-55 mV), all spikes excluded. Red: VSD signal computed with thresholded $V_m$ (-55 mV), only spikes in somatic compartments excluded. **c**, Comparison of VSD signal contributions by bAP (green) and sAP (red), with firing rate overlay (black). **d**, L6 and L5 mean firing rates during time window in dashed box in **c**.

### Fig. 5: Anticorrelation lag time between VSDI and population firing rate for evoked responses

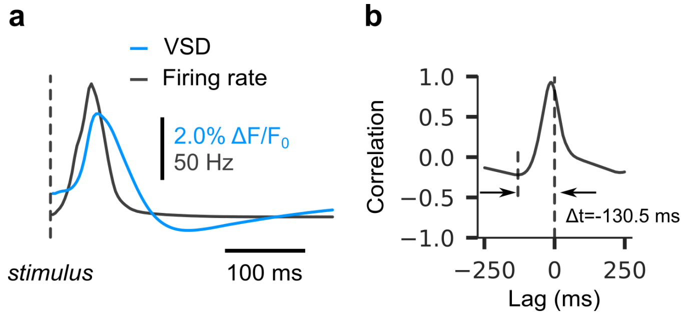


**a**, VSDI activity (blue line) and population firing rate (black line) in a 350 ms poststimulus time window (averaged over n=10 trials). Dashed horizontal line indicates stimulus onset. **b**, Cross-correlogram of traces in **a** (VSD vs. firing rate) for lags spanning the interval [-250 ms, 250 ms]. Time lag associated with anticorrelation peak (-130.5 ms) framed by horizontal arrows.

### Table 1 Wavefront Propagation Velocities

| ***Publication*** | ***Min. speed*** | ***Max. speed*** | ***Wavefront quantification*** | ***Experiment protocol*** | ***Anesthesia*** | ***Brain region*** | ***Animal*** |
|---|---|---|---|---|---|---|---|
| Petersen et al., 2003a | 33 μm/ms (barrel arc) | 60 μm/ms (barrel row) | Gaussian fit (cross-sectional) | whisker deflection (in vivo) | urethane or halothane | barrel cortex | rat P21-P28 |
| Fehérvári et al., 2015 | 47+/-12 μm/ms | 66+/-15 μm/ms | amplitude threshold (50% peak) | 50 μA current injection (in vivo) | urethane | visual cortex (V1) | mouse P56-P140 |
| Ferezou et al., 2006 | 27+/-7 μm/ms (urethane) | 30+/-7 μm/ms (awake) | amplitude threshold (50% peak) | spontaneous (in vivo) | urethane or isoflurane or none | barrel cortex | mouse |
| Lippert et al., 2007 | 200+/-100 μm/ms | 200+/-100 μm/ms | -- | whisker deflection (in vivo) | isoflurane | barrel cortex | rat |
| Petersen et al., 2003b | <10 μm/ms | >100 μm/ms | amplitude threshold (50% peak) | spontaneous (in vivo) | urethane or ketamine/ xylazine or halothane | barrel cortex | rat/mouse P21-P35 |
| Contreras and Llinas, 2001 | 181+/-44 μm/ms (L2/3) | 217+/-53 μm/ms (L5/6) | -- | white matter stimulation 1-5V, 100μs (in vitro) | sodium pentobarbital | visual and somatosens ory cortex | guinea pig |
| Chavane et al., 2011 | 90 μm/ms | 90 μm/ms | 2D Gaussian fit | sinusoidal luminance gratings (in vivo) | althesin (3 mg/kg/h) and pancuronium bromide (0.2 mg/kg/h) | visual cortex | cat |
| Civillico and Contreras, 2005 | 30 μm/ms (single whisker) | 196 μm/ms (multiple whiskers) | amplitude threshold (2x SD of baseline per pixel) | whisker deflection (in vivo) | ketamine-xylazine (100 mg/kg i.p., 20 mg/kg i.p. respectively) | barrel cortex | mouse |
| **NMC** | **~-10 μm/ms** | **~20 μm/ms** | **2D Gaussian fit** | **whisker deflection (in silico)** | **--** | **somato-sensory cortex** | **rat** |

### Algorithm 1: 2D Gaussian surface fit

**input** : A surface arr of size $n \times m$
**output** : A 2D Gaussian fit

```
/* compute 2D surface                                        */
Function 2DGauss(h, x, y, σ_x, σ_y):
    return h · exp −1/2 [ (x/σ_x)^2 + (y/σ_y)^2 ]

/* compute first-order moments of arr                        */
Function Moments(arr):
    total ← sum of entries in arr
    x_0 ← (1/total) ∑ ({row of arr} × {row index})       // 1st moment in x
    y_0 ← (1/total) ∑ ({col of arr} × {col index})       // 1st moment in y
    r ← row of arr at index int(x)
    c ← column of arr at index int(y)
    σ_x ← sqrt( ∑ |(r − index of r) × r|^2 / ∑ r )
    σ_y ← sqrt( ∑ |(c − index of c) × c|^2 / ∑ c )
    h ← max of arr
    return x_0, y_0, h, σ_x, σ_y

/* compute 2D surface                                        */
Function Fitgauss(arr):
    x_0, y_0, h, σ_x, σ_y ← Moments(arr)
    x ← n × m matrix of row indices
    y ← n × m matrix of column indices
    def err_fn (x, y, arr):
        return 2DGauss(x-x_0, y-y_0, h, σ_x, σ_y) - arr
    return LeastSq(err_fn, x, y, arr)        // least-squares fit using
        scipy.optimize

/* main routine                                              */
Function Main(arr):
    params = Fitgauss(arr)
```
